# Transcriptomic analysis reveals hub genes and pathways in response to acetic acid stress in *Kluyveromyces marxianus* during high-temperature ethanol fermentation

**DOI:** 10.1101/2022.06.10.495650

**Authors:** Yumeng Li, Shiqi Hou, Ziwei Ren, Shaojie Fu, Sunhaoyu Wang, Mingpeng Chen, Yan Dang, Hongshen Li, Shizhong Li, Pengsong Li

## Abstract

The thermotolerant yeast *Kluyveromyces marxianus* has great potential for high-temperature ethanol fermentation, but it produces excess acetic acid during high-temperature fermentation, which inhibits ethanol production. The mechanisms of *K. marxianus*’s responses to acetic acid have not been fully understood. In this study, the transcriptomic changes of *K. marxianus* DMKU3-1042 resulted from acetic acid stress during high-temperature ethanol fermentation were investigated based on high-throughput RNA sequencing. We identified 611 differentially expressed genes (DEGs) under acetic acid stress (fold change > 2 or < 0.5, *P*-adjust <0.05), with 166 up-regulated and 445 down-regulated. GO terms and pathways enriched in these DEGs were identified. Protein-protein interaction (PPI) networks were constructed based on the interactions between proteins coded by the DEGs, and hub genes and key modules in the PPI networks were identified. The results in this study indicated that during high-temperature fermentation, acetic acid stress promoted protein catabolism but repressed protein synthesis, which affected the growth and metabolism of *K. marxianus* and led to the decrease of ethanol production. The findings in this study provide a better understanding of the response mechanism of *K. marxianus* to acetic acid stress, and provide a basis for subsequent increase of ethanol production by *K. marxianus*.

## 1. Introduction

In recent years, biofuels have become an important choice of energy substitution all over the world. Among various biofuels, bioethanol is one of the most widely used. The substitution of fossil fuels by bioethanol can effectively reduce the emissions of both air pollutants [1, 2] and CO_2_ [3]. Bioethanol has been produced from sucrose- and starch-rich crops (known as the 1^st^ generation bioethanol). However, these feedstocks conflict with food and feed production. As an alternative to the 1^st^ generation bioethanol, cellulosic ethanol (known as the 2^nd^ generation bioethanol), which utilizes lignocellulose from forestry and agricultural residues, has been developed. To produce cellulosic ethanol, lignocellulose-rich feedstocks are usually subjected to pretreatment (deconstruction of lignocellulose into cellulose, hemicellulose and lignin), saccharification (breakdown of cellulose and hemicellulose into hexose and pentose) and fermentation (co-fermentation of hexose and pentose to ethanol). Although *Saccharomyces cerevisiae* has been widely employed for industrial ethanol fermentation, it lacks genes for pentose utilization, which limits its application in production of cellulosic ethanol. Thermotolerant yeast *Kluyveromyces marxianus* has a number of advantages over *S. cerevisiae* such as thermotolerance, high growth rate and a broad substrate spectrum [4]. Thanks to the thermotolerance of *K. marxianus*, its application in high-temperature fermentation for bioethanol production can help reduce cooling cost, reduce the risk of contamination and achieve simultaneous saccharification and fermentation more efficiently [5]. *K. marxianus* is able to ferment both hexose and pentose without being genetically modified. These characteristics make *K. marxianus* a suitable candidate for cellulosic ethanol production [6].

Various fermentation inhibitors such as weak acids, furan aldehydes and phenolic compounds that are generated during pretreatment of lignocellulosic feedstocks can inhibit subsequent microbial growth, metabolism and ethanol production [7]. Acetic acid is one of the main fermentation inhibitors generated during acid-catalyzed hydrolysis of lignocellulose [8] and has been found to affect the growth and metabolism of *K. marxianus* [9, 10]. In addition, our previous studies found that *K. marxianus* produced more acetic acid during high-temperature fermentation than at lower temperatures, and therefore ethanol fermentation arrested before glucose was fully consumed [11, 12]. Hence, acetic acid is one of the main factors limiting high-temperature ethanol fermentation of *K. marxianus*. Nevertheless, the transcriptomic responses of *K. marxianus* to acetic acid during high-temperature ethanol fermentation have not been fully understood.

The development of high-throughput sequencing technology makes it possible to deeply explore the response mechanisms of yeasts to acetic acid stress. At present, most studies mainly focus on acetic acid stress on *Saccharomyces cerevisiae*, while there are many research gaps in the response and tolerance mechanism of *K. marxianus* to acetic acid stress. Therefore, revealing the response mechanism of acetic acid stress is of great significance to improve the tolerance of *K. marxianus* to acetic acid stress and promote its application in high-temperature ethanol fermentation.

By analyzing the transcriptome changes of *K. marxianus* under acetic acid stress, the response mechanisms of *K. marxianus* to acetic acid stress was revealed in this study. The findings in this study will provide a scientific basis for the construction of acetic acid-tolerant *K. marxianus* strains.

## 2. Materials and methods

### 2.1. Strain, media and culture conditions

*K. marxianus* DMKU3-1042 [5], which was purchased from NITE Biological Resource Center with the deposit number of NBRC 104275, was used throughout this study. YPD medium (10 g/L yeast extract, 20 g/L peptone and 20 g/L glucose) was used for pre-culture of the yeast. After overnight pre-culture in flasks with shaking at 45°C, yeast cells were washed with sterilized water and inoculated into 100-mL serum bottles with 30 mL fermentation medium (10 g/L yeast extract, 20 g/L peptone, 80 g/L glucose and 40 g/L xylose) in each bottle. The initial optical density at 600 nm (OD_600_) of yeast cells in each bottle was set as ∼3.0. To investigate the effect of acetic acid on ethanol fermentation, the concentrations of acetic acid in the fermentation media were set as 0%, 0.25% and 0.3% (w/v), respectively. All fermentation experiments were conducted at 45°C with three biological replicates.

### 2.2. Quantitative analyses of substrates and extracellular metabolites

Broth samples were collected at intervals throughout the fermentation process. The samples were centrifuged at 10,000×g for 1 min. Then the supernatants were diluted by 0.05 mol/L H_2_SO_4_ for 20 times and filtered through filters with 0.45-µm pores. The cell pellets were flash-frozen in liquid nitrogen and stored at −80LJ for subsequent analysis. The concentrations of glucose, xylose, acetic acid, ethanol and glycerol in the fermentation broth were measured by a high-performance liquid chromatography (HPLC) system equipped with an RID-20A refractive index detector (Shimadzu, Japan) and an Aminex HPX-87H column (Bio-Rad, Hercules, CA, USA). The mobile phase was 0.05 mol/L H_2_SO_4_ with a flow rate of 0.6 mL/min. The column temperature and detector temperature were both set as 40°C.

The fermentation parameters were calculated as follows:

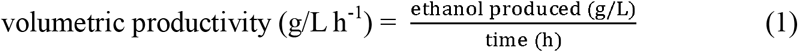

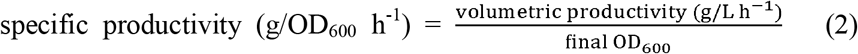

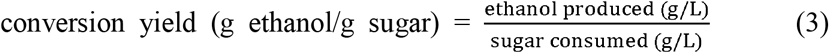

### 2.3. High-throughput RNA sequencing and bioinformatic analysis

To investigate the transcriptomic responses of the yeast to acetic acid, yeast cells collected at the 2nd hour of high-temperature fermentation under the condition of 0.25% acetic acid (treatment group) and no acetic acid (control group) were subjected to high-throughput RNA sequencing. Total RNA samples were extracted from the cell pellets using the EZNA Yeast RNA Kit (Omega Bio-tek, Doraville, CA, USA) and then sent to Shanghai Majorbio Bio-pharm Technology Co., Ltd. (Shanghai, China) for quality and quantity evaluation, cDNA library construction and high-throughput sequencing. The genome sequence of *K. marxianus* DMKU3-1042 in the NCBI database (accession number: PRJDA65233) [13] was used as the reference genome. After removing the adaptors and the low-quality reads, the clean reads were aligned to the reference genome using HISAT2 [14]. The differentially expressed genes (DEGs) were identified using DESeq2 [15]. The resulting *P* values were adjusted using the Benjamin and Hochberg’s approach for controlling the false discovery rate. Genes with adjusted *P* (*P*-adjust) values less than 0.05 found by DESeq2 were considered as differentially expressed. Gene Ontology (GO) enrichment analysis of the DEGs was performed using the GOseq R package [16]. KOBAS software was used for KEGG pathway enrichment analysis [17]. GO terms and KEGG pathways with *P-*adjust values less than 0.05 were considered significantly enriched.

### 2.4. Protein-protein interaction (PPI) network analysis

The STRING database (http://string-db.org) [18] was used to construct PPI networks of the identified DEGs. Given that the PPI information of *K. marxianus* was not included in the STRING database, we chose *Kluyveromyces lactis* as the reference. Then the PPI network data was imported into the Cytoscape software for subsequent analysis [19]. The centrality parameters (degree, betweenness, eigenvector) were analyzed using CentiScaPe 2.2 [20]. Nodes with higher centrality values than average were identified as hub nodes. The major PPI networks were constructed based on the intersection of the hub nodes identified based on the three selected centrality parameters. The most significant modules in a major PPI network were identified using Molecular Complex Detection (MCODE) plugin [21] with a K-score value of 5. The hub genes in a PPI network were ranked based on the MCC algorithm in CytoHubba plugin of Cytoscape [22].

## 3. Results and discussion

### 3.1. Acetic acid repressed high-temperature ethanol fermentation of *K. marxianus*

The cell concentration was monitored throughout the high-temperature ethanol fermentation process. When no acetic acid was added, the OD_600_ reached a peak of ∼9.4 within 8 hours (Fig. 1a). When acetic acid was added to the fermentation media, however, the growth of *K. marxianus* was significantly repressed, with maximum OD_600_ of ∼4.3 and ∼4.0 appeared at 8 h in the groups treated with 0.25% and 0.3% acetic acid, respectively (Fig. 1a).

**Fig. 1.**
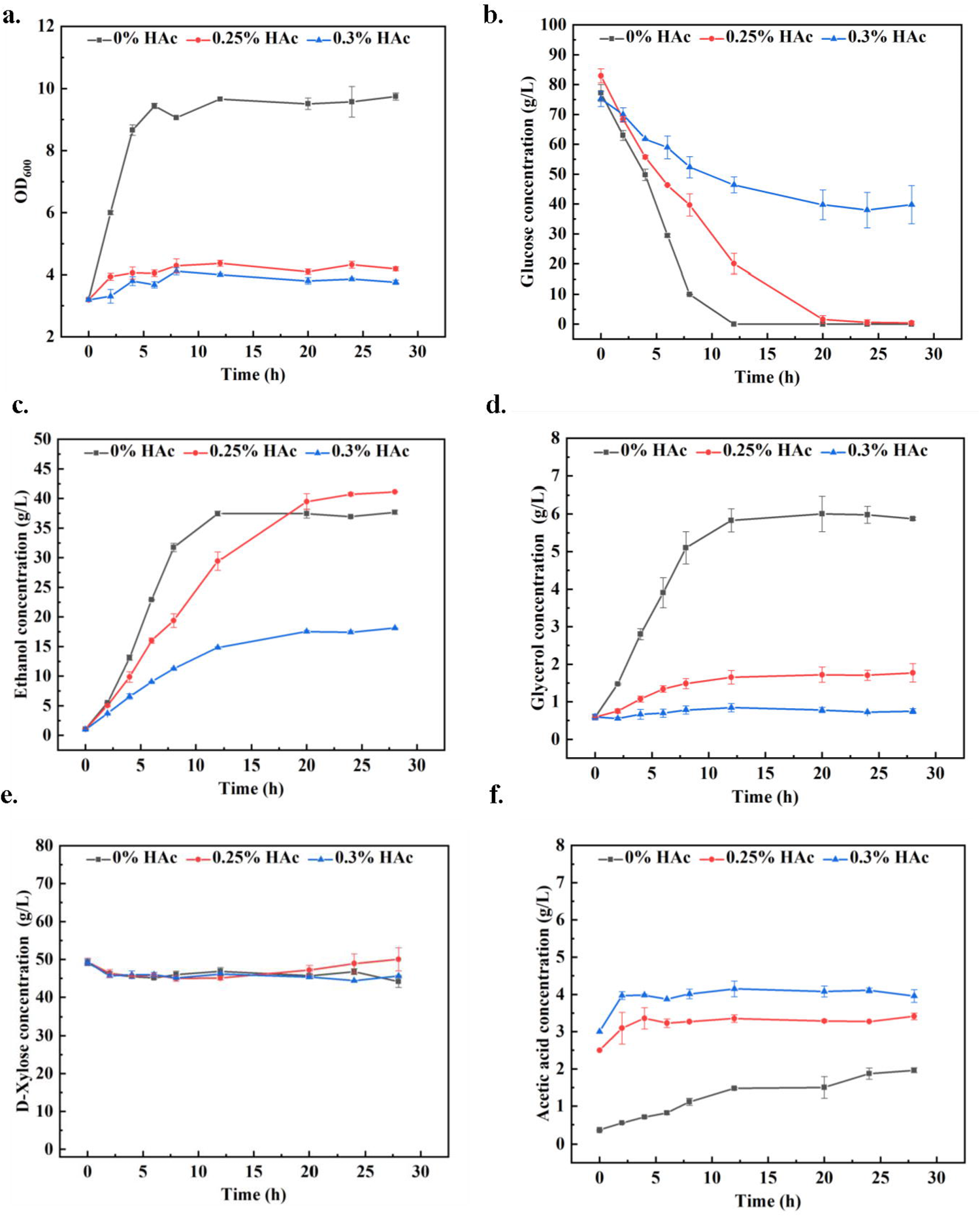
Changes in (a) OD_600_, concentrations of (b) glucose, (c) ethanol, (d) glycerol, (e) xylose and (f) acetic acid during fermentation at 45°C under different concentrations of acetic acid.

According to the HPLC results, the metabolism of *K. marxianus* was also inhibited by acetic acid. Glucose consumption in the control group was the fastest, with the glucose consumed completely within 10 hours, while the glucose in the group treated with 0.25% acetic acid was completely consumed within more than 20 hours (Fig. 1b). Glucose consumption in the group treated with 0.3% acetic acid was not only the slowest, but it also stopped after 20 h, with around 40 g/L glucose remained unconsumed (Fig. 1b). Acetic acid treatment also slowed down ethanol generation. The ethanol concentration in the control group reached a peak of 37 g/L within 12 h, while the ethanol concentrations in the groups treated with 0.25% and 0.3% acetic acid reached corresponding maximum values of 40 and 18 g/L at around 24 h, respectively (Fig. 1c). Interestingly, total glycerol production of the group treated by 0.25% acetic acid was less than that of the control group (Fig. 1d), making the conversion yield of the former (0.49 g ethanol/g sugar) higher than that of the latter (0.45 g ethanol/g sugar) (Table 1). To our surprise, although treatment with 0.25% acetic acid inhibited cell growth and slowed down ethanol generation, the specific productivity in this group was significantly higher than the that in the control group (Table 1).

**Table 1.** Fermentation results (0-24 h) in this study.

| Fermentation parameters | 0% HAc | 0.25% HAc | 0.3% HAc |
| --- | --- | --- | --- |
| Initial OD<sub>600</sub> | 3.19 | 3.19 | 3.19 |
| Final OD<sub>600</sub> | 9.74 | 4.19 | 3.76 |
| Volumetric productivity (g/L h<sup>-1</sup>) | 1.31 | 1.43 | 0.61 |
| Specific productivity (g/OD<sub>600</sub> h<sup>-1</sup>) | 0.13 | 0.34 | 0.16 |
| Conversion yield (g ethanol/g sugar) | 0.45 | 0.49 | 0.44 |

### 3.2. Descriptive statistics of RNA-Seq data

In order to reveal the transcriptomic responses of *K. marxianus* induced by acetic acid, the cells in the group treated with 0.25% acetic acid and the control group were sampled at 2 h, and then subjected to total RNA extraction. According to the results of RNA quality evaluation (Table S1), the RNA samples were qualified for library construction. After library construction and paired-end sequencing, 297.7 million paired-end raw reads were generated for the six samples in total, ranging from 42.3 to 56.9 million reads per sample. After removing the low complexity reads, adaptor, primer sequences and rRNA sequences, 41.7-56.3 million clean reads were yielded for each sample (Table S2). All the raw data were deposited to the China National GeneBank database (CNGBdb) under the accession number of XXXXX. Then we employed HISAT2 to align the clean reads to the reference genome, and the alignment results show that 92.97-95.07% of clean reads for each sample were uniquely mapped to the reference genome (Table S3). We also performed PCA to investigate if samples with the same treatment cluster together. According to the PCA result, the first two principal components explained more than 89% of the variability among the samples, and acetic acid treated samples and control samples were grouped in different clusters (Fig. S1). This result indicated that the transcriptome profiles was significantly changed after acetic acid treatment. The acetic acid treated samples fell in the negative direction of the PC1 axis, while the control samples fell in the positive direction. In PC2, one sample in the control group did not cluster with others.

### 3.3. Identification of DEGs and functional enrichment analysis

According to differential expression analysis, 611 DEGs (fold change > 2 or < 0.5, *P*-adjust < 0.05) were identified in the samples treated with 0.25% acetic acid compared with the control group, with 166 up-regulated and 455 down-regulated (Fig. 2a). Among the up-regulated DEGs, those with fold changes > 10 accounted for 18.07 %, those with fold changes between 5 and 10 accounted for 35.54%, and those with fold changes between 2 and 5 accounted for 46.39%. Among the down-regulated DEGs, those with fold changes < 0.1 accounted for 4.49%, those with fold changes between 0.1 and 0.2 accounted for 8.31%, and those with fold changes between 0.2 and 0.5 accounted for 87.19%.

**Fig. 2.**
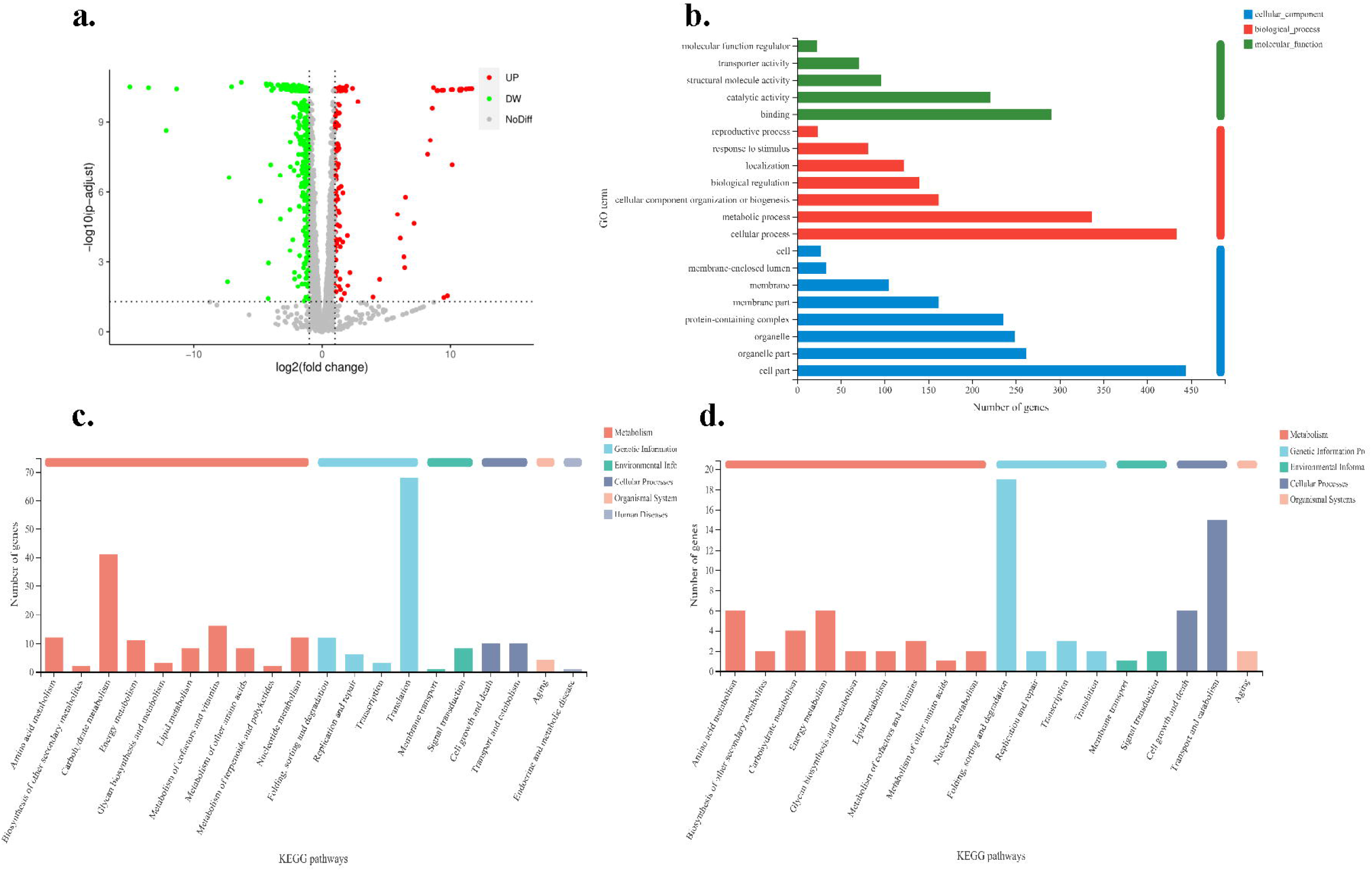
Identification of differentially expressed genes (DEGs) induced by acetic acid. (a) volcano plot of the DEGs; (b) GO annotation result of the DEGs; (c) KEGG pathway annotation result of down-regulated DEGs; (d) KEGG pathway annotation result of up-regulated DEGs.

The DEGs were functionally categorized into GO functional classes. The top 5 GO terms with the largest number of DEGs were cell part (GO:0044464), cellular process (GO:0009987), metabolic process (GO:0008152), binding (GO:0005488) and organelle part (GO:0044422) (Fig. 2b). The DEGs were also mapped to the pathways in the KEGG database. The top 5 pathways with the most down-regulated DEGs were translation, carbohydrate metabolism, metabolism of cofactors and vitamins, amino acid metabolism and nucleotide metabolism and folding, sorting and degradation (Fig. 2c), while the top 5 pathways with the most up-regulated DEGs were folding, sorting and degradation, transport and catabolism, amino acid metabolism, energy metabolism and cell growth and death (Fig. 2d).

To further analyze the functions of the DEGs induced by acetic acid, GO and KEGG enrichment analyses were conducted. According to the result of GO enrichment analysis, the significantly enriched GO terms in the DEGs were related to nucleotide phosphorylation (such as GO:0046939, GO:0006165 and GO:0006757), protein synthesis (such as GO:0022625 and GO:0015934) and carbon metabolism (such as GO:0006096 and GO:0006006), indicating that acetic acid may act on mitochondria, ribosome-related molecular functions, cellular components, biological processes, etc. to influence the energy conversion process in cells and thus the growth and metabolism of microorganisms (Fig. 3a and Table S4). The result of KEGG pathway enrichment show that ribosome (map03010), fructose and mannose metabolism (map00051), glycolysis/gluconeogenesis (map00010), proteasome (map03050), etc. were enriched in the DEGs (Fig. 3b and Table S5).

**Fig. 3.**
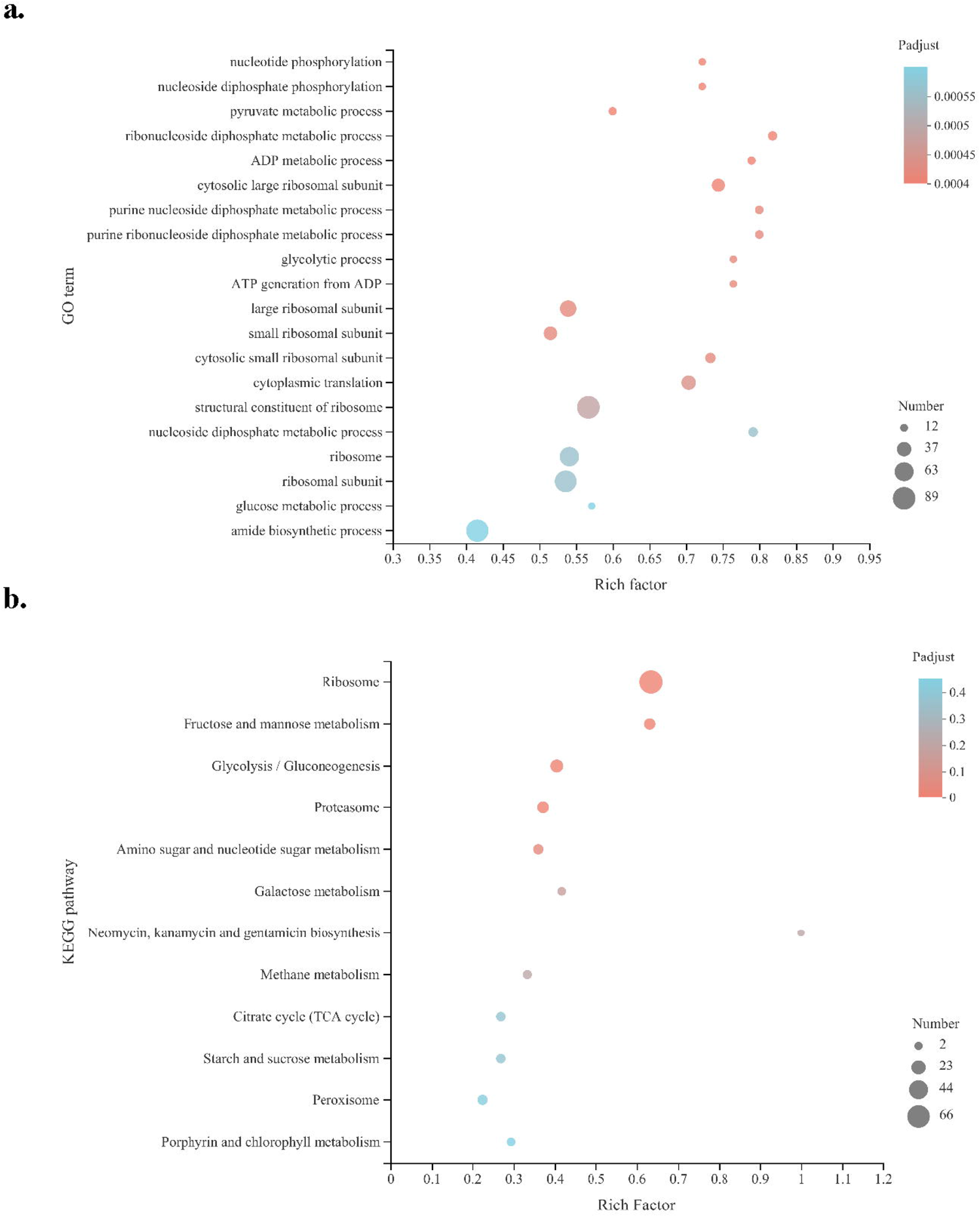
Enrichment analyses of the DEGs in this study. (a) GO terms enriched in the DEGs; (b) KEGG pathways enriched in the DEGs. Rich factor is the ratio of DEG number annotated in this GO term (or KEGG pathway) to all gene number annotated in this GO term (or KEGG pathway). Greater rich factor means greater effect of acetic acid on the analyzed GO term (or KEGG pathway).

### 3.4 PPI network analysis

In order to reveal functional interactions among the proteins coding by the DEGs, PPI networks of all DEGs, up-regulated DEGs and down-regulated DEGs were constructed, respectively. Four hundred and seventeen proteins associated with the DEGs were matched with the STRING database and used to construct the PPI networks. The threshold of interaction score was set to > 0.4 and the unconnected nodes were hidden. The constructed PPI networks of all DEGs consisted of 329 nodes and 2041 edges (Fig. 4), while those of up-regulated DEGs and down-regulated DEGs consisted of 73 nodes, 116 edges (Fig. S2), and 225 nodes, 1612 edges (Fig. S3), respectively.

**Fig. 4.**
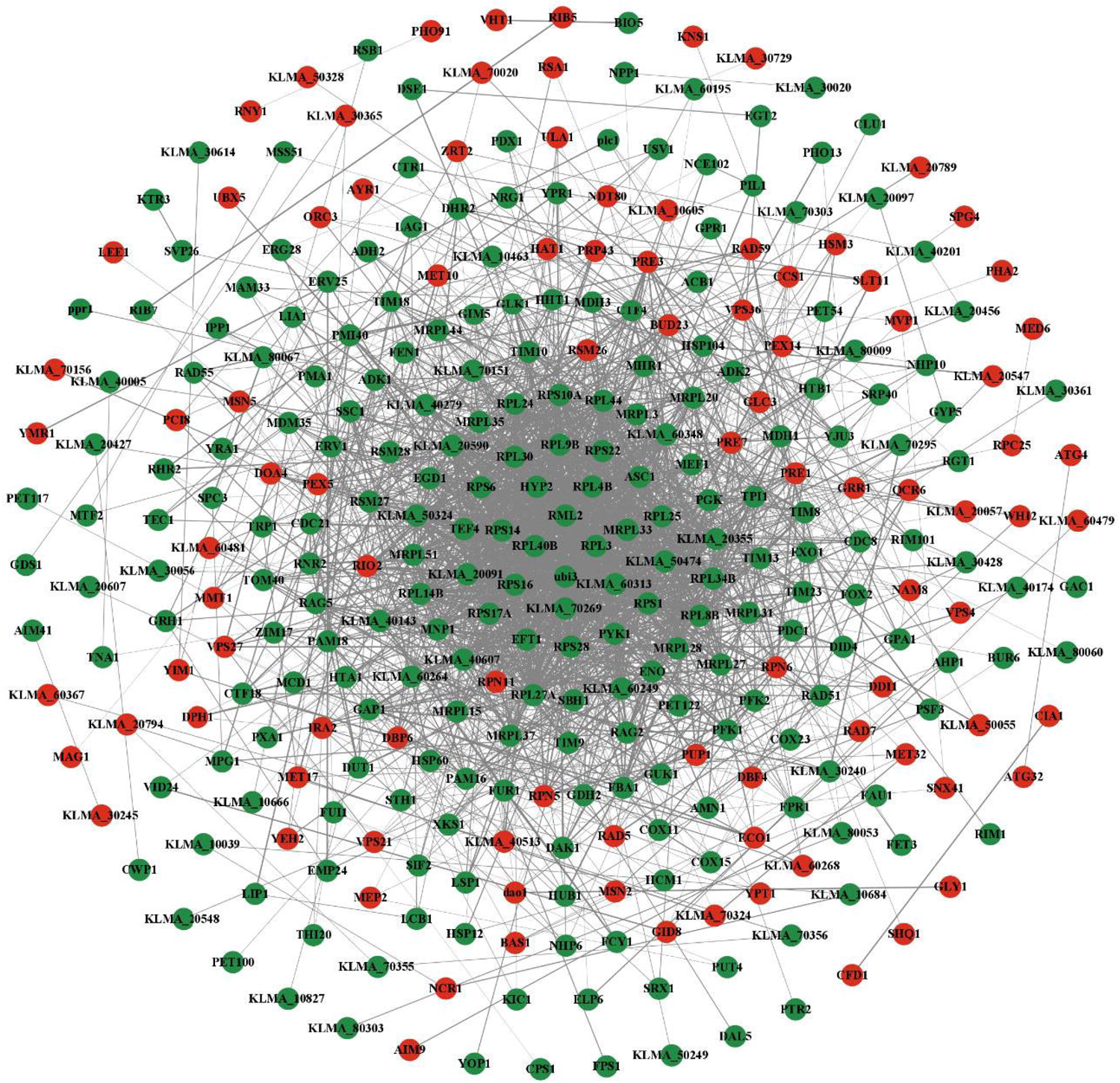
Protein-protein interaction (PPI) network of the proteins coded by the DEGs in this study. Red nodes represent proteins coded by up-regulated DEGs; green nodes represent proteins coded by down-regulated DEGs; edges represent protein-protein interactions, and thicker edges indicate stronger interactions.

To obtain the major PPI network of up-regulated DEGs, the topological connectivity of each node was determined based on the centrality parameters degree, betweenness and eigenvector. According to the analyses of centrality parameters, twenty-two, eighteen and sixteen nodes were identified from the PPI network of up-regulated DEGs with the values of degree, betweenness and eigenvector above average, respectively. Seven nodes showed all these three centrality parameters above average and formed the major PPI network of up-regulated DEGs with 12 edges, which was equivalent to 24.1% of the PPI network of up-regulate DEGs (Fig. 5a and Fig. S4). These hub genes were ranked by MCC value (Fig. 5b and Table S6). A single module was identified from this major PPI network (Fig. 5c). Similarly, the major PPI network of down-regulated DEGs with 32 nodes and 316 edges was obtained (Fig. 6a and Fig. S5). The top 10 hub genes with higher MCC values were identified and sequentially ordered (Fig. 6b and Table S7). In addition, three significant modules were identified via the MCODE plugin (Fig. 6c).

**Fig. 5.**
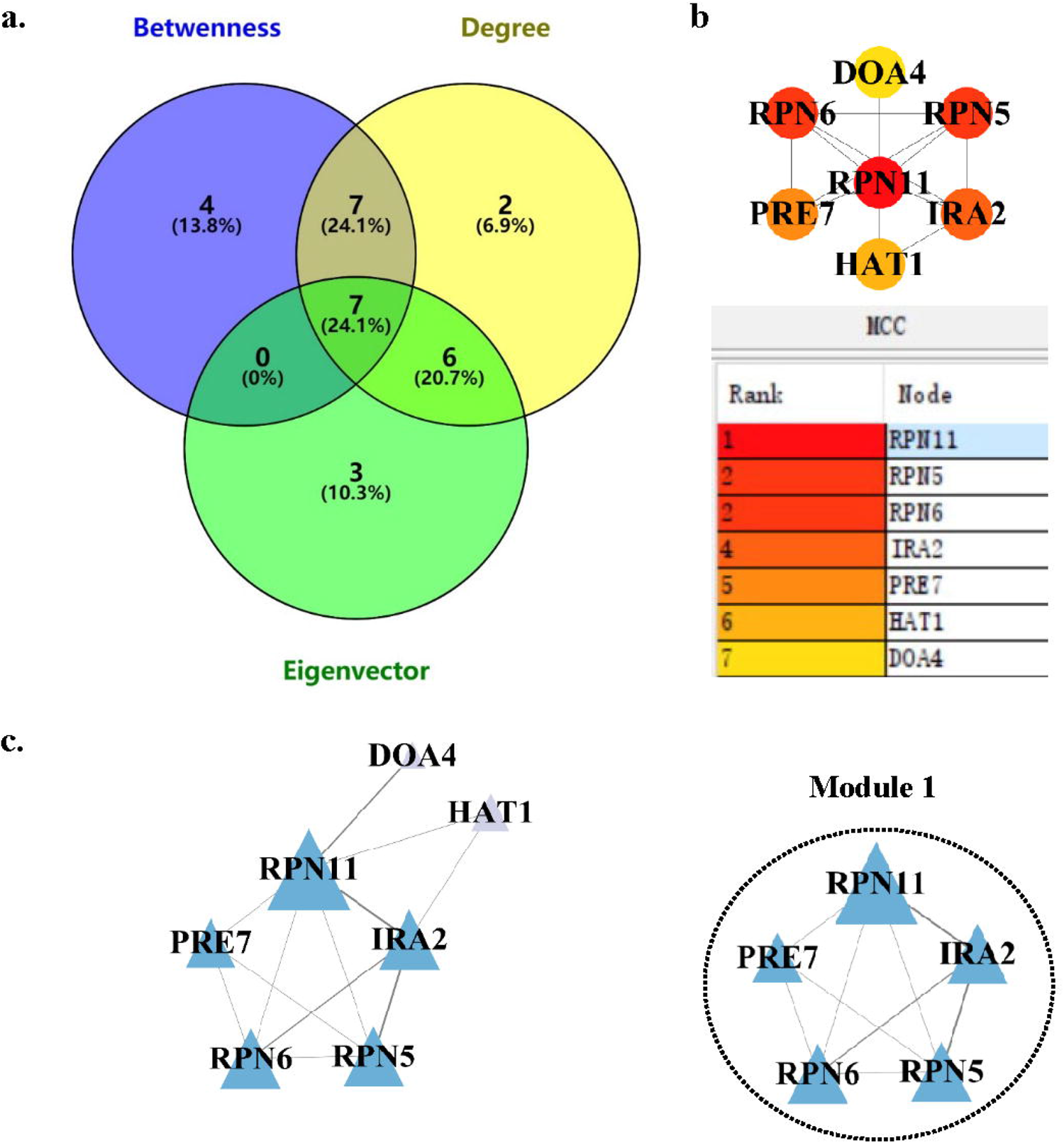
Major PPI networks of up-regulated DEGs and module analysis. (a) Venn diagram showing nodes with values of centrality parameters (degree, Betweenness, and eigenvector) above average. (b) Major PPI networks of up-regulated DEGs and ranking of the up-regulated hub genes. Node color reflects the degree of connectivity, the redder a node, the greater its MCC value. (c) MCODE analysis. Module 1: score = 4.5.

**Fig. 6.**
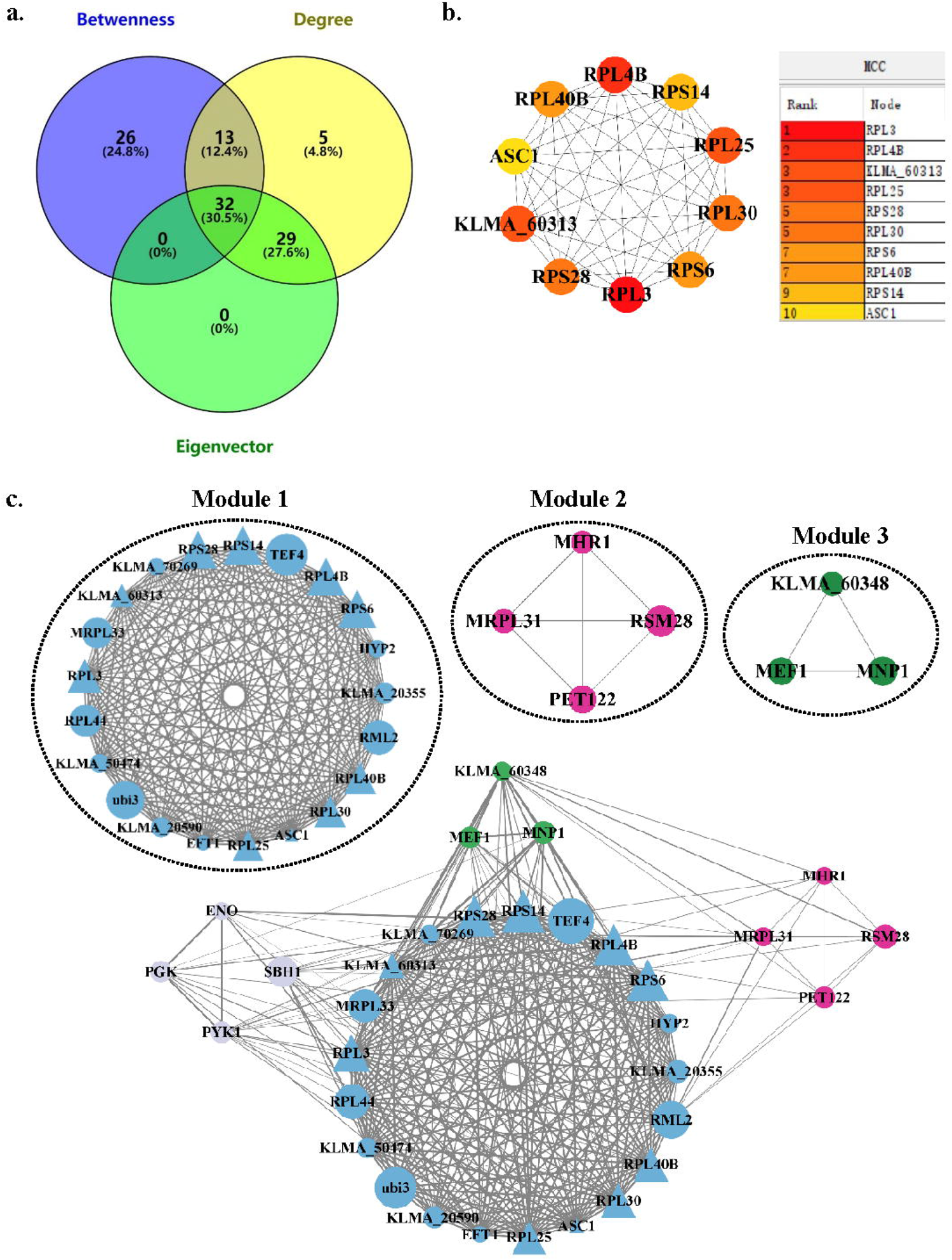
Major PPI networks of down-regulated DEGs and module analysis. (a) Venn diagram showing nodes with values of centrality parameters (degree, Betweenness, and eigenvector) above average. (b) Top 10 down-regulated hub genes with highest MCC values in the major PPI network of down-regulated DEGs. Node color reflects the degree of connectivity, the redder a node, the greater its MCC value. (c) MCODE analysis. Module 1: score = 20.8; Module 2: score = 4; Module 3: score = 3. Triangles represent hub genes; the sizes of the circles indicate the MCC values.

To further investigate the functions of genes in the identified modules, we performed GO and KEGG enrichment analyses for these genes. We found that GO terms and KEGG pathways related to proteasome were enriched in the only identified module of major PPI network of up-regulated DEGs (Tables S8-S9). For the major PPI network of down-regulated DEGs, GO terms associated with ribosome, mitochondrial ribosome and mitochondrial translation were enriched in modules 1, 2 and 3, respectively, and an KEGG pathway related to ribosome was also enriched in module 1, but no KEGG pathway was enriched in modules 2 and 3 due to the limited numbers of nodes in these modules. These results indicated that acetic acid stress promoted protein catabolism but repressed protein synthesis, which affected the growth and metabolism of *K. marxianus* and led to the decrease of ethanol production.

## 4. Conclusions

In this study, the transcriptomic changes of *K. marxianus* DMKU3-1042 resulted from acetic acid stress during high-temperature ethanol fermentation were investigated based on high-throughput RNA sequencing. Hub genes and key modules in the PPI networks of the DEGs were identified. The results indicated that during high-temperature fermentation, acetic acid stress promoted protein catabolism but repressed protein synthesis, which affected the growth and metabolism of *K. marxianus* and led to the decrease of ethanol production. The findings in this study provide a better understanding of the response mechanism of *K. marxianus* to acetic acid stress, and provide a basis for subsequent increase of ethanol production by *K. marxianus*.

## Acknowledgements

This work was supported by National Key Research and Development Program of China (2021YFC3200602), National Undergraduate Training Program for Innovation and Entrepreneurship (202110022074, 202198039), and Beijing Municipal Education Commission through the Innovative Transdisciplinary Program “Ecological Restoration Engineering” (GJJXK210102).

## References

[1] Goldemberg J, Ethanol for a sustainable energy future, Science 315 (2007) 808–810.

[2] Salvo A, Brito J, Artaxo P, Geiger FM, Reduced ultrafine particle levels in Sao Paulo’s atmosphere during shifts from gasoline to ethanol use, Nat Commun 8 (2017) 77.

[3] Scully MJ, Norris GA, Falconi TMA, Macintosh DL, Carbon intensity of corn ethanol in the United States: state of the science, Environmental Research Letters 16 (2021).

[4] Fonseca GG, Heinzle E, Wittmann C, Gombert AK, The yeast Kluyveromyces marxianus and its biotechnological potential, Appl Microbiol Biotechnol 79 (2008) 339–54.

[5] Limtong S, Sringiew C, Yongmanitchai W, Production of fuel ethanol at high temperature from sugar cane juice by a newly isolated Kluyveromyces marxianus, Bioresour Technol 98 (2007) 3367–3374.

[6] Nonklang S, Abdel-Banat BM, Cha-Aim K, Moonjai N, Hoshida H, Limtong S, Yamada M, Akada R, High-temperature ethanol fermentation and transformation with linear DNA in the thermotolerant yeast Kluyveromyces marxianus DMKU3-1042, Appl Environ Microbiol 74 (2008) 7514–7521.

[7] Wang D, Wu D, Yang X, Hong J, Transcriptomic analysis of thermotolerant yeast Kluyveromyces marxianus in multiple inhibitors tolerance, RSC Adv 8 (2018) 14177–14192.

[8] An J, Kwon H, Kim E, Lee YM, Ko HJ, Park H, Choi I-G, Kim S, Kim KH, Kim W, Choi W, Tolerance to acetic acid is improved by mutations of the TATA-binding protein gene, Environ Microbiol 17 (2015) 656–669.

[9] Rugthaworn P, Murata Y, Machida M, Apiwatanapiwat W, Hirooka A, Thanapase W, Dangjarean H, Ushiwaka S, Morimitsu K, Kosugi A, Arai T, Vaithanomsat P, Growth inhibition of thermotolerant yeast, Kluyveromyces marxianus, in hydrolysates from cassava pulp, Appl Biochem Biotechnol 173 (2014) 1197–1208.

[10] Martynova J, Kokina A, Kibilds J, Liepins J, Scerbaka R, Vigants A, Effects of acetate on Kluyveromyces marxianus DSM 5422 growth and metabolism, Appl Microbiol Biotechnol 100 (2016) 4585–4594.

[11] Fu X, Li P, Zhang L, Li S, Understanding the stress responses of Kluyveromyces marxianus after an arrest during high-temperature ethanol fermentation based on integration of RNA-Seq and metabolite data, Appl Microbiol Biotechnol 103 (2019) 2715–2729.

[12] Li P, Tan X, Fu X, Dang Y, Li S, Metabolomic analysis reveals Kluyveromyces marxianus’s stress responses during high-temperature ethanol fermentation, Process Biochem 102 (2021) 386–392.

[13] Lertwattanasakul N, Kosaka T, Hosoyama A, Suzuki Y, Rodrussamee N, Matsutani M, Murata M, Fujimoto N, Suprayogi Tsuchikane K, Limtong S, Fujita N, Yamada M, Genetic basis of the highly efficient yeast Kluyveromyces marxianus: complete genome sequence and transcriptome analyses, Biotechnol Biofuels 8 (2015) 47.

[14] Kim D, Langmead B, Salzberg SL, HISAT: a fast spliced aligner with low memory requirements, Nat Methods 12 (2015) 357–360.

[15] Love MI, Huber W, Anders S, Moderated estimation of fold change and dispersion for RNA-seq data with DESeq2, Genome Biol 15 (2014) 550.

[16] Young MD, Wakefield MJ, Smyth GK, Oshlack A, Gene ontology analysis for RNA-seq: accounting for selection bias, Genome Biol 11 (2010) R14.

[17] Mao X, Cai T, Olyarchuk JG, Wei L, Automated genome annotation and pathway identification using the KEGG Orthology (KO) as a controlled vocabulary, Bioinformatics 21 (2005) 3787–93.

[18] Szklarczyk D, Gable AL, Nastou KC, Lyon D, Kirsch R, Pyysalo S, Doncheva NT, Legeay M, Fang T, Bork P, Jensen LJ, Von Mering C, The STRING database in 2021: customizable protein-protein networks, and functional characterization of user-uploaded gene/measurement sets, Nucleic Acids Res 49 (2021) D605–D612.

[19] Shannon P, Markiel A, Ozier O, Baliga NS, Wang JT, Ramage D, Amin N, Schwikowski B, Ideker T, Cytoscape: a software environment for integrated models of biomolecular interaction networks, Genome Res 13 (2003) 2498–2504.

[20] Scardoni G, Petterlini M, Laudanna C, Analyzing biological network parameters with CentiScaPe, Bioinformatics 25 (2009) 2857–9.

[21] Bader GD, Hogue CW, An automated method for finding molecular complexes in large protein interaction networks, BMC Bioinf 4 (2003) 2.

[22] Chin CH, Chen SH, Wu HH, Ho CW, Ko MT, Lin CY, cytoHubba: identifying hub objects and sub-networks from complex interactome, BMC Syst Biol 8 Suppl 4 (2014) S11.

