## supplementary material for "Transcriptomic analysis reveals hub genes and pathways in response to acetic acid stress in *Kluyveromyces marxianus* during high-temperature ethanol fermentation"

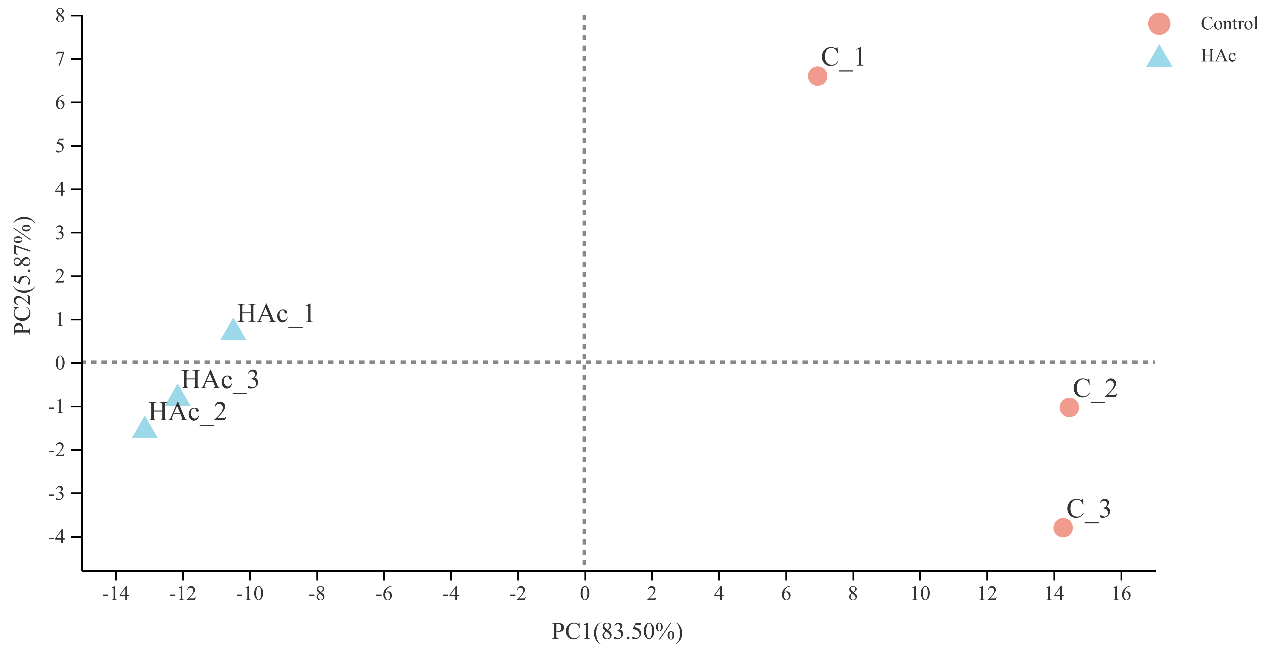

**Fig. S1** PCA scatter plot of gene expression.

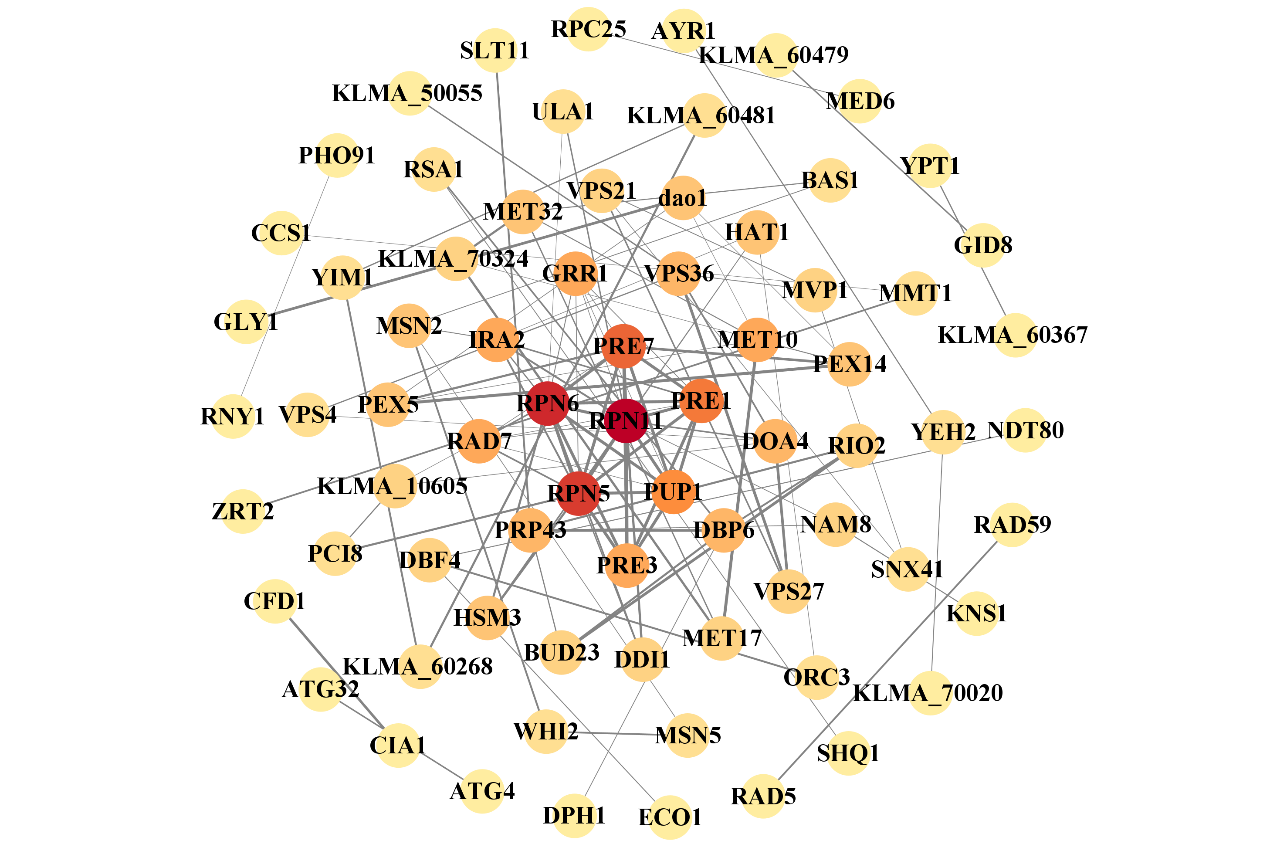

**Fig. S2** PPI network of up-regulated DEGs.

Nodes represent proteins, and the darker color of a node, the higher its MCC value; edges represent protein-protein interactions, and thicker edges indicate stronger interactions.

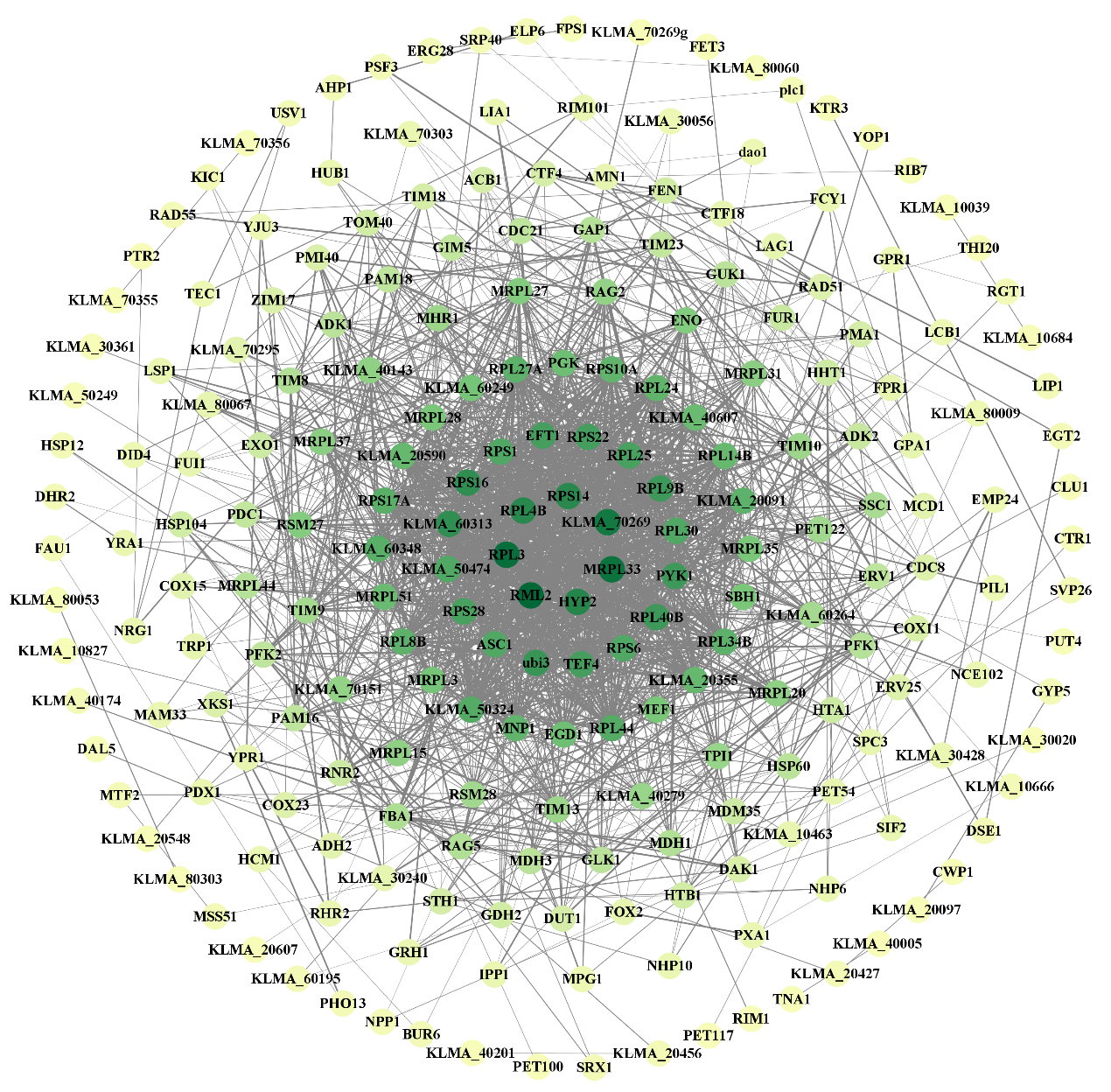

**Fig. S3** PPI network of down-regulated DEGs.

Nodes represent proteins, and the darker color of a node, the higher its MCC value; edges represent protein-protein interactions, and thicker edges indicate stronger interactions.

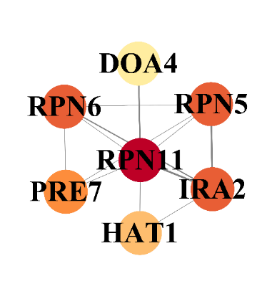

**Fig. S4** Major PPI network of up-regulated DEGs.

Nodes represent proteins, and the darker color of a node, the higher its MCC value; edges represent protein-protein interactions, and thicker edges indicate stronger interactions.

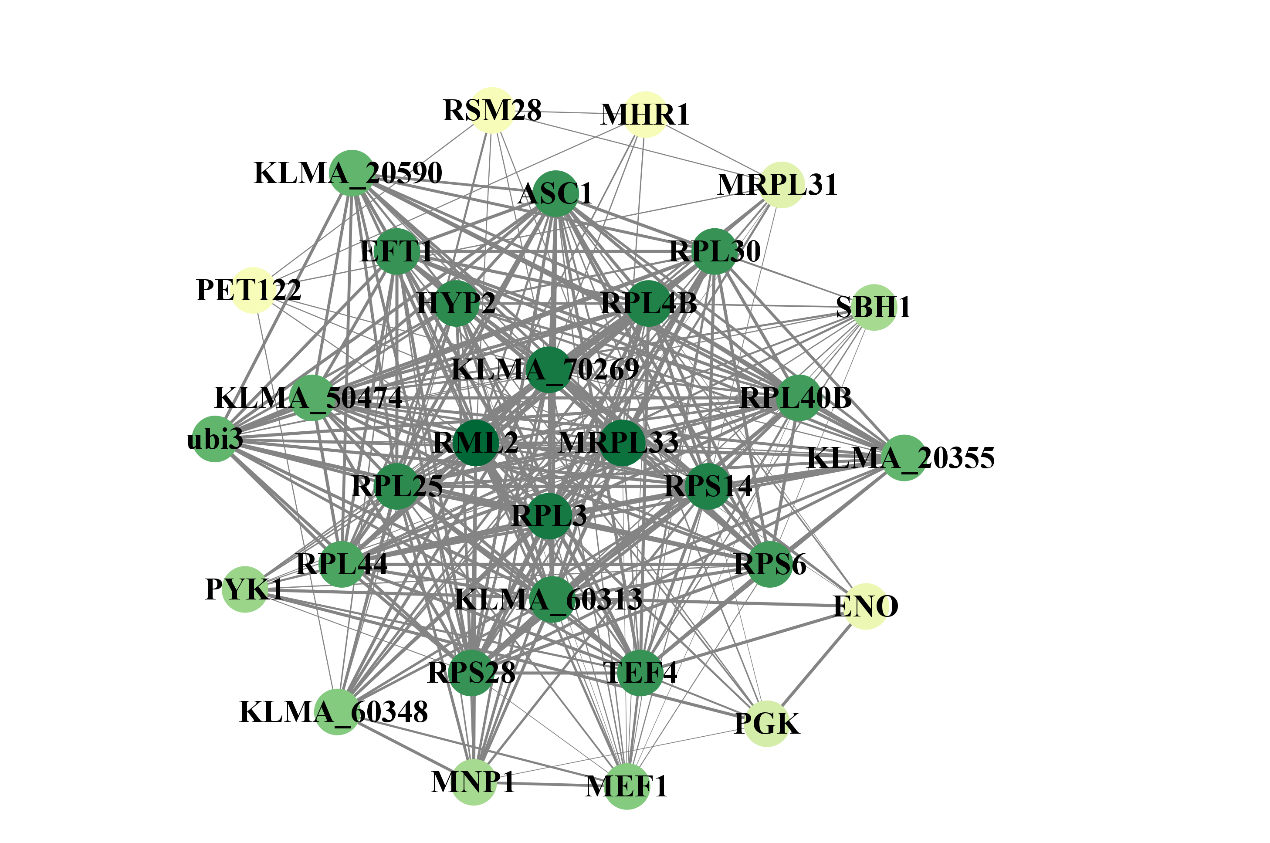

**Fig. S5** Major PPI network of down-regulated DEGs.

Nodes represent proteins, and the darker color of a node, the higher its MCC value; edges represent protein-protein interactions, and thicker edges indicate stronger interactions.

**Table S1** Quality evaluation of RNA samples in this study.

| Group | Sample | Concentration  (ng/μL) | Total (μg) | OD_260_/OD_280_ | OD_260_/OD_230_ | RIN |
| --- | --- | --- | --- | --- | --- | --- |
| HAc | HAc_1 | 474.40 | 16.60 | 2.09 | 2.38 | 10.00 |
| HAc | HAc_2 | 378.40 | 13.24 | 2.13 | 2.46 | 10.00 |
| HAc | HAc_3 | 262.70 | 9.19 | 2.15 | 2.44 | 10.00 |
| Control | C_1 | 629.90 | 22.05 | 2.18 | 2.41 | 10.00 |
| Control | C_2 | 317.90 | 11.13 | 2.16 | 2.47 | 10.00 |
| Control | C_3 | 658.90 | 23.06 | 2.17 | 2.47 | 10.00 |

**Table S2** Raw and clean sequence details in this study.

| Sample | Raw reads | Raw bases | Clean reads | Clean bases | Error rate  (%) | Q20 (%) | Q30 (%) | GC content (%) |
| --- | --- | --- | --- | --- | --- | --- | --- | --- |
| HAc_1 | 56935702 | 8597291002 | 56325880 | 8347943232 | 0.0255 | 97.74 | 93.75 | 43.95 |
| HAc_2 | 52817164 | 7975391764 | 52225956 | 7709684741 | 0.0256 | 97.71 | 93.68 | 43.85 |
| HAc_3 | 52330720 | 7901938720 | 51707486 | 7676640027 | 0.0256 | 97.71 | 93.68 | 43.92 |
| C_1 | 45203110 | 6825669610 | 44605648 | 6628821827 | 0.0258 | 97.63 | 93.52 | 44.6 |
| C_2 | 42250302 | 6379795602 | 41668990 | 6178210169 | 0.0258 | 97.61 | 93.52 | 45.01 |
| C_3 | 48144372 | 7269800172 | 47502238 | 7029631209 | 0.0260 | 97.54 | 93.34 | 44.71 |

**Table S3** Descriptive statistics of mapping rate.

| Sample | Total reads | Total mapped  reads | Multiple mapped reads | Uniquely mapped reads |
| --- | --- | --- | --- | --- |
| HAc_1 | 56325880 | 54771301(97.24%) | 1638088(2.91%) | 53133213(94.33%) |
| HAc_2 | 52225956 | 50713277(97.10%) | 2160631(4.14%) | 48552646(92.97%) |
| HAc_3 | 51707486 | 50203925(97.09%) | 1948796(3.77%) | 48255129(93.32%) |
| C_1 | 44605648 | 43365077(97.22%) | 959319(2.15%) | 42405758(95.07%) |
| C_2 | 41668990 | 40509797(97.22%) | 944407(2.27%) | 39565390(94.95%) |
| C_3 | 47502238 | 46138043(97.13%) | 1256449(2.65%) | 44881594(94.48%) |

**Table S4** GO enrichment result.

| Ranking | Description | Term Type | Number |
| --- | --- | --- | --- |
| 1 | nucleotide phosphorylation | BP | 13 |
| 2 | nucleoside diphosphate phosphorylation | BP | 13 |
| 3 | pyruvate metabolic process | BP | 15 |
| 4 | ribonucleoside diphosphate metabolic process | BP | 18 |
| 5 | ADP metabolic process | BP | 15 |
| 6 | cytosolic large ribosomal subunit | CC | 32 |
| 7 | purine nucleoside diphosphate metabolic process | BP | 16 |
| 8 | purine ribonucleoside diphosphate metabolic process | BP | 16 |
| 9 | glycolytic process | BP | 13 |
| 10 | ATP generation from ADP | BP | 13 |
| 11 | large ribosomal subunit | CC | 48 |
| 12 | small ribosomal subunit | CC | 34 |
| 13 | cytosolic small ribosomal subunit | CC | 22 |
| 14 | cytoplasmic translation | BP | 38 |
| 15 | structural constituent of ribosome | MF | 89 |
| 16 | nucleoside diphosphate metabolic process | BP | 19 |
| 17 | ribosome | CC | 66 |
| 18 | ribosomal subunit | CC | 82 |
| 19 | glucose metabolic process | BP | 12 |
| 20 | nucleotide phosphorylation | BP | 13 |

**Table S5** KEGG enrichment result.

| Ranking | Description | First Category | Number |
| --- | --- | --- | --- |
| 1 | Ribosome | Genetic Information Processing | 66 |
| 2 | Fructose and mannose metabolism | Metabolism | 12 |
| 3 | Glycolysis / Gluconeogenesis | Metabolism | 17 |
| 4 | Proteasome | Genetic Information Processing | 13 |
| 5 | Amino sugar and nucleotide sugar metabolism | Metabolism | 9 |
| 6 | Galactose metabolism | Metabolism | 5 |
| 7 | Neomycin, kanamycin and gentamicin biosynthesis | Metabolism | 2 |
| 8 | Methane metabolism | Metabolism | 6 |
| 9 | Citrate cycle (TCA cycle) | Metabolism | 7 |
| 10 | Starch and sucrose metabolism | Metabolism | 7 |
| 11 | Peroxisome | Cellular Processes | 9 |
| 12 | Porphyrin and chlorophyll metabolism | Metabolism | 5 |

**Table S6** Information of hub nodes in PPI network of up-regulated DEGs.

| Hub nodes | Protein names | Function |
| --- | --- | --- |
| *RPN11* | 26S proteasome regulatory subunit RPN11 | isopeptidase activity, Lys63-specific deubiquitinase activity Lys63, metallopeptidase activity, metallopeptidase activity, peroxisome fission, proteasome assembly, proteasome-mediated ubiquitin-dependent protein catabolic process, proteasome storage granule assembly. |
| *RPN5* | 26S proteasome regulatory subunit RPN5 | proteasome-mediated ubiquitin-dependent protein catabolic process, protein deneddylation. |
| *RPN6* | 26S proteasome regulatory subunit RPN6 | structural molecule activity, proteasome assembly, proteasome-mediated ubiquitin-dependent protein catabolic process. |
| *IRA2* | inhibitory regulator protein IRA2 | regulation of GTPase activity GTP, signal transduction. |
| *PRE7* | Proteasome component C5 | threonine-type endopeptidase activity, proteasomal ubiquitin-independent protein catabolic process, proteasome-mediated ubiquitin-dependent protein catabolic process. |
| *HAT1* | Histone acetyltransferase type B catalytic subunit | chromatin binding; H4 histone acetyltransferase activity; histone binding; cellular response to DNA damage stimulus; subtelomeric heterochromatin assembly. |
| *DOA4* | Ubiquitin carboxyl-terminal hydrolase | protein deubiquitination; regulation of DNA replication; ubiquitin-dependent protein catabolic process via the multivesicular body sorting pathway. |

**Table S7** Information of hub nodes in PPI network of down-regulated DEGs.

| Hub nodes | Protein names | Function |
| --- | --- | --- |
| *RPL3* | 60S ribosomal protein L3 | structural constituent of ribosome, maintenance of translational fidelity, ribosomal large subunit assembly, rRNA processing. |
| *RPL4B* | 60S ribosomal protein L4-B | structural constituent of ribosome, translation. |
| KLMA_60313 | 40S ribosomal protein S20 | structural constituent of ribosome, maturation of SSU-rRNA from tricistronic rRNA transcript (SSU-rRNA, 5.8S rRNA, LSU-rRNA), translation. |
| *RPL25* | Ribosomal protein L23 | RNA binding, structural constituent of ribosome, ribosomal large subunit assembly, translation. |
| *RPS28* | 40S ribosomal protein S28 | structural constituent of ribosome, positive regulation of nuclear-transcribed mRNA catabolic process, deadenylation-dependent decay, translation. |
| *RPL30* | 60S ribosomal protein L30 | pre-mRNA 5'-splice site binding, structural constituent of ribosome, negative regulation of mRNA splicing, via spliceosome, rRNA processing. |
| *RPS6* | 40S ribosomal protein S6 | structural constituent of ribosome, translation. |
| *RPL40B* | Ubiquitin-60S ribosomal protein L40 | structural constituent of ribosome, translation. |
| *RPS14* | 40S ribosomal protein S14 | structural constituent of ribosome, translation. |
| *ASC1* | Guanine nucleotide-binding protein subunit beta-like | adaptor/regulatory modules in signal transduction, pre-mRNA processing, cytoskeleton assembly. |

**Table S8** GO terms enriched in the genes in the modules of major PPI networks in this study.

| **up/down** | **module** | **Description** | **Term Type** | **Number** |
| --- | --- | --- | --- | --- |
| up | module 1 | Proteasome storage granule | CC | 4 |
|  |  | Proteasome regulatory particle, lid subcomplex | CC | 3 |
|  |  | Ubiquitin-dependent protein catabolic process | BP | 4 |
| down | module 1 | Ribosome | CC | 19 |
|  |  | Translation | BP | 20 |
|  |  | Structural constituent of ribosome | MF | 17 |
|  |  | Ribosomal subunit | CC | 16 |
|  |  | Cytosolic ribosome | CC | 14 |
|  |  | Large ribosomal subunit | CC | 11 |
|  |  | Cytosolic large ribosomal subunit | CC | 9 |
|  |  | Cytoplasmic translation | BP | 7 |
|  |  | Translational elongation | BP | 5 |
|  |  | Small ribosomal subunit | CC | 5 |
|  |  | Cytosolic small ribosomal subunit | CC | 4 |
|  |  | Translation elongation factor activity | MF | 3 |
|  |  | Ribosome assembly | BP | 4 |
|  |  | Ribosome biogenesis | BP | 7 |
| down | module 2 | Mitochondrial ribosome | CC | 4 |
|  |  | Ribosomal subunit | CC | 4 |
|  |  | Mitochondrial protein complex | CC | 4 |
|  |  | Structural constituent of ribosome | MF | 4 |
|  |  | Mitochondrial small ribosomal subunit | CC | 2 |
|  |  | Mitochondrial large ribosomal subunit | CC | 2 |
| down | module 3 | Mitochondrial translation | BP | 3 |

**Table S9** KEGG pathways enriched in the genes in the modules of major PPI networks in this study.

| **up/down** | **module** | **Description** | **First Category** | **Number** |
| --- | --- | --- | --- | --- |
| up | module 1 | Proteasome | Genetic Information Processing | 4 |
| down | module 1 | Ribosome | Genetic Information Processing | 16 |
